# Kinesin-1 transports morphologically distinct intracellular virions during vaccinia infection

**DOI:** 10.1101/2022.04.15.488437

**Authors:** Amadeus Xu, Angika Basant, Sibylle Schleich, Timothy P Newsome, Michael Way

**Affiliations:** Cellular signalling and cytoskeletal function laboratory, The Francis Crick Institute, 1 Midland Road, London, NW1 1AT, UK; London Research Institute, Cancer Research UK, 44 Lincoln’s Inn Fields, London, WC2A 3PX, UK; German Cancer Research Center (DKFZ), 69120, Heidelberg, Germany; School of Life and Environmental Sciences, The University of Sydney, Sydney, New South Wales, Australia; Department of Infectious Disease, Imperial College, London W2 1PG, UK

## Abstract

Intracellular mature virions (IMV) are the first and most abundant infectious form of vaccinia virus to assemble during its replication cycle. IMV can undergo microtubule-based motility, but their directionality and the motor involved in their transport remain unknown. Here, we demonstrate that IMV, like intracellular enveloped virions (IEV), the second form of vaccinia, undergo anterograde transport and recruit kinesin-1. In vitro reconstitution of virion transport reveals that IMV and IEV move toward microtubule plus-ends with respective velocities of 0.66 and 0.56 μm/s. Quantitative imaging establishes IMV and IEV recruit an average of 65 and 115 kinesin-1 motor complexes respectively. In the absence of kinesin-1 there is a near-complete loss of in vitro motility and defects in the cellular spread of both virions. Our observations demonstrate kinesin-1 transports two morphologically distinct forms of vaccinia. Reconstitution of vaccinia-based microtubule motility in vitro provides a new model to investigate how motor number and regulation impacts transport of a bona fide kinesin-1 cargo.

## Introduction

Kinesin-1, the founding member of the kinesin superfamily, anchors and transports a diverse range of cellular cargoes, including vesicles, organelles, protein complexes, and RNPs towards the plus end of microtubules (Hirokawa et al., 2009; Verhey and Hammond, 2009). Kinesin-1 is a heterotetramer consisting of two heavy chains that each contain an N-terminal motor domain that is necessary for movement, and two light chains which play important roles in motor regulation and cargo binding (Bloom et al., 1988; Hackney and Stock, 2000; Kaan et al., 2011; Vale et al., 1985). In humans the kinesin heavy chain (KHC) is represented by three different genes that encode closely related isoforms, KIF5A, KIF5B and KIF5C. KIF5B appears to be ubiquitously expressed, while KIF5A and KIF5C are neuronal specific (Kanai et al., 2000). Each KIF5 heavy chain homodimer associates near their C-terminus with the heptad repeats of two copies of one of four light chain isoforms (KLC1-4) (Miki et al., 2001). While KLC2 is ubiquitously expressed and KLC1 is found in most cell types, the other isoforms are tissue specific (Junco et al., 2001; Rahman et al., 1998). Despite the importance of kinesin-1 in transport of many cellular cargoes, we lack a thorough understanding of kinesin-1 motor-cargo relationships, including motor activation as well as their number and organization on cargoes. This is in part due to the lack of well-defined exemplary kinesin-1 cargoes, and the challenge of detecting kinesin-1 on moving cargo using fluorescence-based imaging methods.

Kinesin-1 is also used by a number of different viruses to enhance their replication cycles especially during their egress from infected cells (Diefenbach et al., 2002; Dodding and Way, 2011; DuRaine et al., 2018; Jouvenet et al., 2004; Pegg et al., 2021; Rietdorf et al., 2001; Strunze et al., 2011). Understanding how viruses recruit kinesin-1 via a limited set of proteins offers a great opportunity to understand the molecular basis of motor recruitment and regulation, as well as their organization on a defined cargo. We previously demonstrated that during vaccinia virus infection, intracellular enveloped virions (IEV) recruit kinesin-1 to mediate their microtubule-dependent transport from their perinuclear site of assembly to the plasma membrane (Rietdorf et al., 2001). Disruption of the ability of IEV to recruit kinesin-1 leads to a dramatic reduction in viral transport to the plasma membrane and cell-to-cell spread of the virus (Rietdorf et al., 2001; Ward and Moss, 2001). Kinesin-1 is recruited to IEV by the interaction of A36, an integral IEV membrane protein with the tetratricopeptide repeats (TPR) of the kinesin light chain (Ward and Moss, 2004). A36 interacts with the TPR via a bipartite tryptophan acidic motif that is also found in many other cellular proteins that bind kinesin-1 (Dodding et al., 2011; Pernigo et al., 2013). More recently, the viral E2/F12 complex that associates with IEV moving on microtubules (Dodding et al., 2009), was shown to enhance kinesin-1 binding to A36, suggesting the virus also regulates motor recruitment (Carpentier et al., 2015; Gao et al., 2017).

However, in infected cells, IEV only comprise a small proportion of total cytoplasmic virions compared to its precursor the intracellular mature virion (IMV), the first infectious form of vaccinia virus assembled during infection (Carpentier et al., 2017; Leite and Way, 2015; Payne and Kristenson, 1979). IEV are formed when IMV acquire an additional membrane cisterna from the trans-Golgi network (TGN) or early endosomes (Leite and Way, 2015; Schmelz et al., 1994; Tooze et al., 1993). This envelopment results in the outer surface of IEV having a very different composition of viral proteins from the IMV including the presence of A36 (Smith et al., 2002). Previous analysis demonstrates that IMV can move at velocities up to 3 μm/s and are susceptible to nocodazole treatment, strongly implicating microtubules in their transport (Sanderson et al., 2000; Ward, 2005). It is thought this motility is important to transport IMV from their site of assembly close to the nucleus towards the TGN to facilitate membrane envelopment and IEV formation (Sanderson et al., 2000; Ward, 2005). In addition, microtubule transport of IMV to the cell periphery may play a role in the cell-to-cell spread of vaccinia as IMV are also capable of directly budding at the plasma membrane (Meiser et al., 2003; Tsutsui, 1983). Curiously, movement of IMV on microtubules has not been imaged directly and the identity of the motor(s) responsible for their translocation to the TGN or plasma membrane remains to be established. Given this, we set out to identify the motor responsible for IMV transport using complementary in vitro and cell-based assays. Our analysis reveals that IMV recruit kinesin-1 albeit at significantly lower levels than IEV. Moreover, kinesin-1 is the major motor driving IMV motility in vitro and its loss leads to a significant defect in virion spread during infection.

## Results and Discussion

### IMV undergo plus end directed microtubule motility

To analyse IMV motility, we infected HeLa cells with a recombinant vaccinia strain ΔB5 lacking the viral protein B5 which is essential for IEV formation (Engelstad and Smith, 1993; Wolffe et al., 1993) that also encodes RFP-tagged core protein A3 for visualisation (Arakawa et al., 2007). Live-imaging of ΔB5 RFP-A3 infected cells labelled with SiR-Tubulin reveals that IMV undergo a variety of movements, including linear transport along microtubules (MTs) and diffusion within the MT network as well as static association with MTs (Fig. 1A and B and Videos 1 and 2). Moreover, disrupting the MT network with nocodazole results in loss of IMV motility in agreement with previous observations (Ward, 2005) (Fig. S1, A-C and Video 3). To analyse IMV movements in detail, we performed automated single particle tracking of fluorescently labelled virions using TrackMate (Fig. 1C) (Tinevez et al., 2017). Periods of active virion transport were discriminated from phases of diffusive and/or confined motion within each trajectory using TraJ (Fig. 1C) (Wagner et al., 2017). Using this approach, quantitative analysis of IMV sub-trajectories undergoing active transport reveals that they move at an average velocity of 0.61 ± 0.35 μm/s over an average run length of 1.72 ± 1.73 μm (Fig. 1D). These values are lower than IEV which move both faster and further, 0.88 ± 0.04 μm/s and 6.44 ± 0.37 μm average velocity and run length respectively (Dodding et al., 2011).

**Figure 1.**
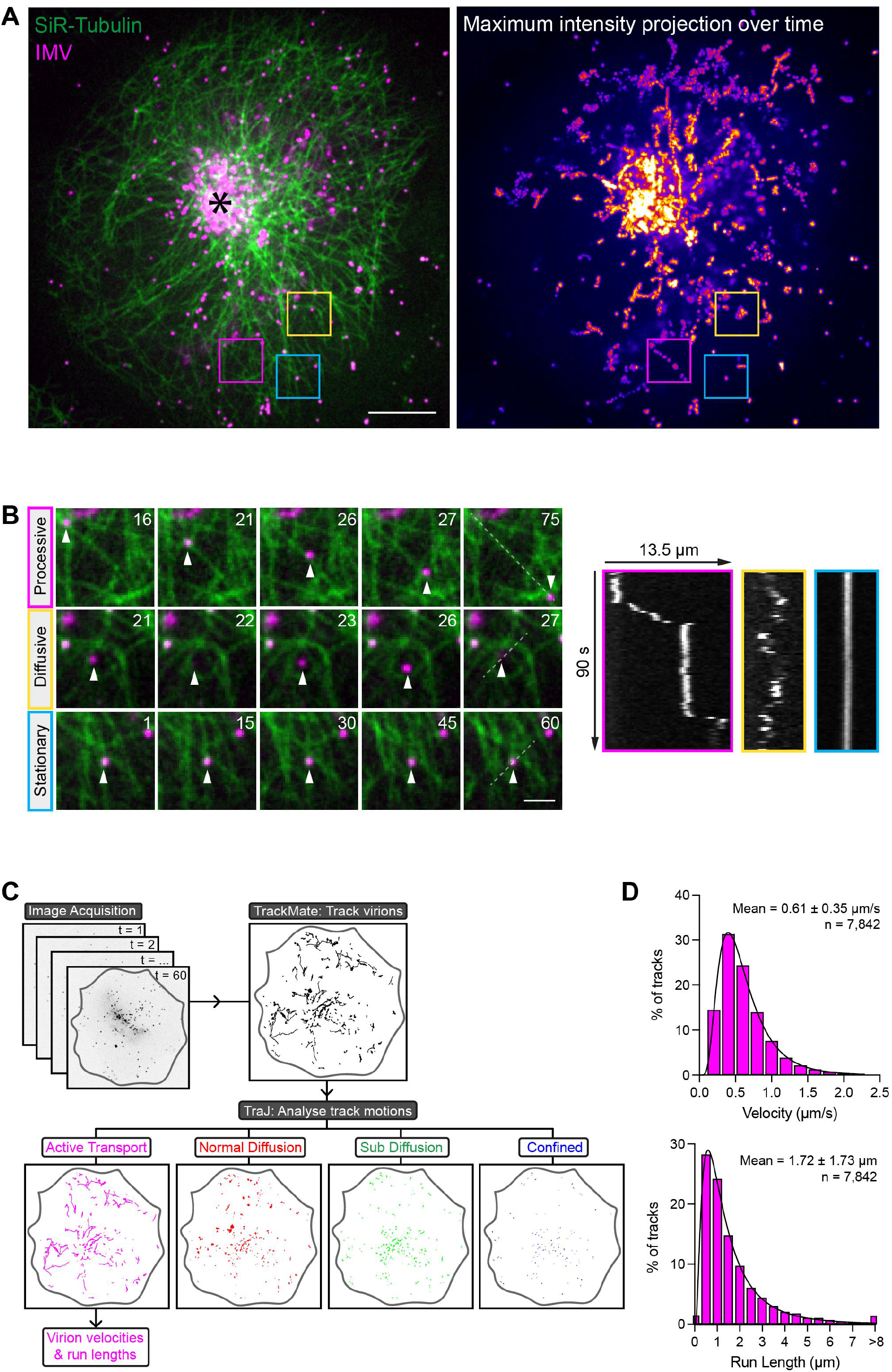
Analysis of microtubule-based motility of IMV in cells. **(A)** Still from time lapse movie showing a HeLa cell labelled with SiR-Tubulin (green) at 7.5 hours post infection with the ΔB5 RFP-A3 virus (magenta) to visualise microtubules and IMV respectively (see Video 1). The asterisk indicates the perinuclear site of IMV assembly. Coloured boxed regions are enlarged in B. Maximum intensity projection of IMV channel over 90 seconds is shown on the right. Scale bar, 20 μm. **(B)** Enlarged boxed regions from A illustrate examples of processive, diffusive and stationary IMV (magenta) movements on microtubules (green) (see Video 2). Time in seconds is indicated in each frame. Corresponding kymographs for each IMV motion over 90 seconds are shown on the right. Scale bar, 2 μm. **(C)** Image acquisition and analysis pipeline used to track virions and categorise their constituent movements as either active motion, normal-/sub-diffusion or confined using Trackmate and TraJ. **(D)** Histograms of the velocities and run lengths of IMV undergoing active motion using automated tracking and analysis. N = 7842 virus runs from 15 ΔB5-infected cells in 3 independent experiments.

The directionality of IMV movements is hard to assess in cells, as the dense microtubule network, especially near the nucleus makes it difficult to determine whether virions are moving on single or bundled microtubules. The typical radial microtubule organisation is also disrupted during vaccinia infection, which also compounds the challenge in determining microtubule polarity and direction of transport (Ploubidou et al., 2000). Reconstitution of microtubule-based motility in vitro has provided major insights into the properties and regulation of kinesin-1 (Block et al., 1990; Chiba et al., 2021; Friedman and Vale, 1999; Hooikaas et al., 2019; Jiang et al., 2019; Seitz and Surrey, 2006; Svoboda et al., 1993). Moreover, microtubule-dependent transport of Herpes Simplex Virus in cell extracts has been reconstituted in vitro (Lee et al., 2006; Wolfstein et al., 2006). Given this and to overcome the issues of microtubule organization in vaccinia infected cells, we established an in vitro assay to analyse IMV motility on purified single microtubules using extracts from ΔB5 RFP-A3 infected HeLa cells (Fig. 2A and B). In parallel, we also analysed the movement of IEV, which are distinguishable from IMV by the presence of A36, using extracts from cells infected with the Western Reserve (WR) strain of vaccinia expressing A36-YdF-YFP RFP-A3 (Fig. 2B). The A36-YdF recombinant virus was used as it is deficient in actin-based motility but not microtubule-based transport (Rietdorf et al., 2001; Ward and Moss, 2001). We observed that both IMV and IEV could move along GMPCPP-stabilised microtubules in an ATP-dependent fashion as the presence of the non-hydrolysable ATP analogue, AMPPNP, inhibited virion motility (Fig. 2C and D and Videos 4 and 5). Consistent with our cell-based observations, IMV moved at an average velocity of 0.66 ± 0.14 μm/s (Fig. 2D). In contrast, IEV moved slightly slower in the in vitro motility assay (0.56 ± 0.08 μm/s). Both viruses had similar run lengths averaging ~8-9 μm, often reaching the microtubule end where they sometimes remain stationary rather than detaching (Fig. 2C and D).

**Figure 2.**
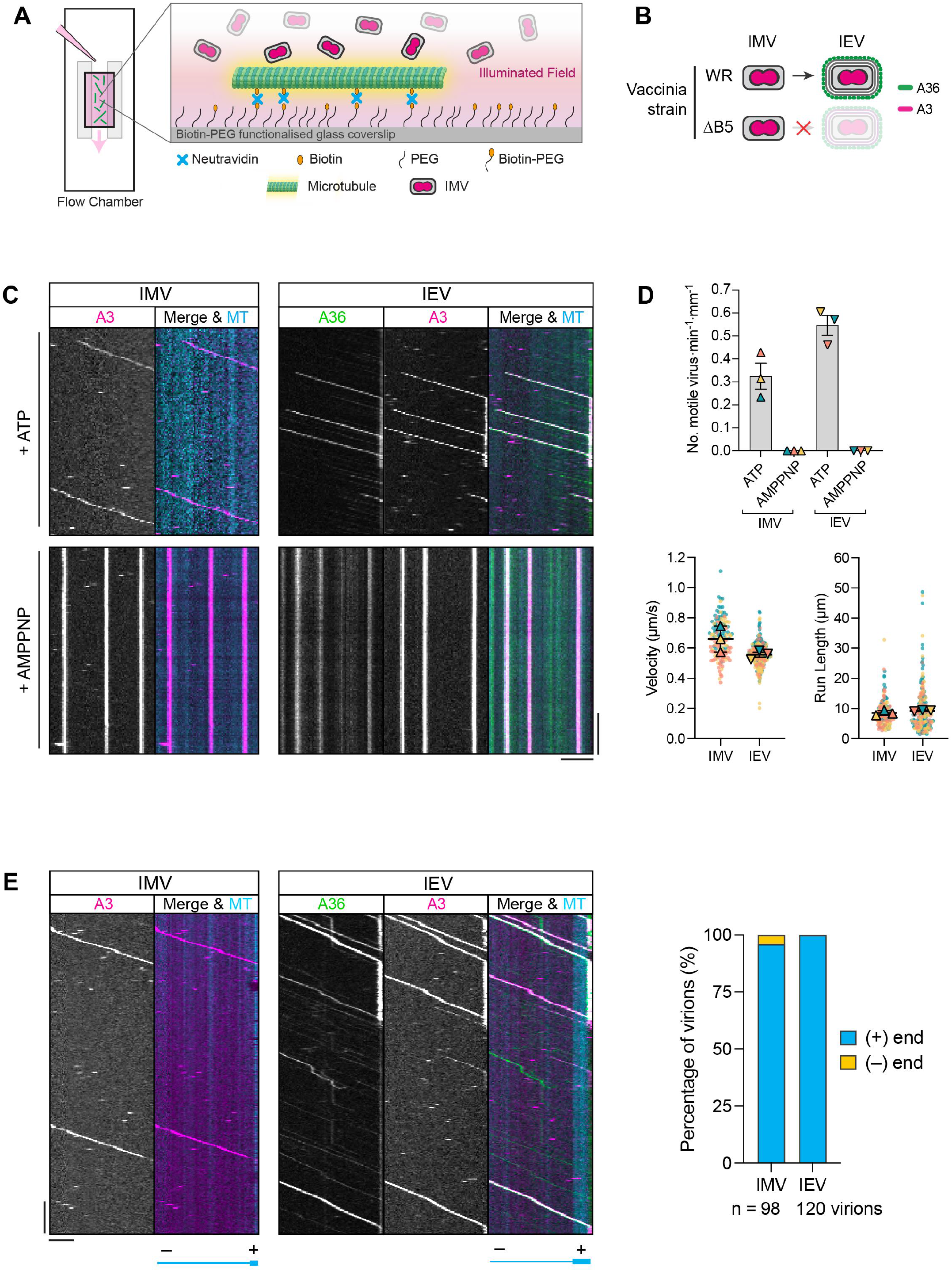
Analysis of microtubule-based IMV and IEV movements in vitro. **(A)** Schematic of in vitro flow chamber illustrating the attachment of biotin- and fluorescently labelled microtubules to biotin-PEG functionalised glass coverslip via a neutravidin link. RFP-tagged IMV are visualised following addition of infected cell extracts into the chamber. **(B)** Schematic of the intracellular virions produced by wild-type Western Reserve (WR) or recombinant ΔB5 strains. Intracellular mature virions (IMV) are labelled by RFP-A3 only while intracellular enveloped virions (IEV) are identified by RFP-A3 and A36-YFP markers. **(C)** Example kymographs of IMV or IEV moving on GMPCPP-microtubules (cyan) in vitro in the presence of either 2 mM ATP or AMPPNP (see Videos 4 and 5). Scale bars, 30 s (vertical) and 5 μm (horizontal). **(D)** SuperPlots showing IMV and IEV in vitro motility rate, virus velocities and run lengths in the presence of ATP or AMPPNP. Error bars represent mean and SEM from 3 independent experiments in which 146 IMV and 263 IEV were analysed. **(E)** Kymographs of IMV or IEV moving on polarity-marked microtubules (cyan) in vitro (See Videos 6 and 7). Microtubule plus (+) and minus (-) ends are indicated below the images. The bar chart (right) shows percentage of IMV and IEV moving towards microtubule (+) or (−) ends. N = 98 (IMV) or 120 (IEV) virions from 3 independent experiments. Scale bars, 30 s (vertical) and 5 μm (horizontal).

Interestingly, IMV and IEV always translocate towards one microtubule end and were never observed moving bidirectionally or travelling in opposite directions on the same microtubule, suggesting that they move exclusively to either the plus- or minus-end. The in vitro unidirectional motility of IEV is likely towards the plus-end given they recruit kinesin-1 in infected cells (Carpentier et al., 2015; Dodding et al., 2011; Rietdorf et al., 2001; Ward and Moss, 2004). In vitro assays using polarity-marked microtubules with bright plus-ends confirmed this is indeed the case (Fig. 2E and Video 6). Interestingly, IMV also move towards the microtubule plus-end in 96% of runs, suggesting their transport is driven by a kinesin (Fig. 2E and Video 7). It is likely that the 4% of IMV moving to the minus-end are false-positives due to mislabelling of microtubule plus-ends resulting from microtubule shearing and reannealing events during their preparation (Fallesen et al., 2017).

### Kinesin-1 is recruited to IMV and IEV

Our observations with polarity marked microtubules prompted us to analyse whether kinesin-1 associates with IMV in infected cells. Immunofluorescence analysis of WR infected HeLa cells reveals that IMV and IEV (identified by the absence or presence of A36 respectively) recruit endogenous KIF5B, KLC1 and KLC2 (Fig. 3A and S1D). Furthermore, endogenous kinesin-1 heavy and light chains as well as GFP-tagged KLC1 and KLC2 associate with IMV in ΔB5-infected cells (Fig. 3B and S1D-F). Strikingly, the fluorescence intensity of endogenous kinesin-1 appeared brighter on IEV compared to IMV (Fig. 3A and S1D). Quantification of heavy and light chain fluorescence intensities on virions in WR infected cells demonstrates IEV recruit ~2-fold more kinesin-1 than IMV, despite the latter comprising the majority of virions assembled during infection (Fig. 3C). This difference in levels suggests that IEV have more binding sites and/or greater affinity for kinesin-1 than IMV. Interestingly, IMV produced in ΔB5-infected cells recruited greater levels of KLC2 compared to IMV in WR-infected cells (Fig. 3C). However, no such difference was observed with KLC1. This may suggest that competition exists between IEV and IMV for binding KLC2/KIF5B tetramers during WR-infection. Curiously, pulldown assays demonstrate that the IEV proteins A36 and E2/F12 have preferential binding for KLC1 and KLC2 isoforms (Gao et al., 2017).

**Figure 3.**
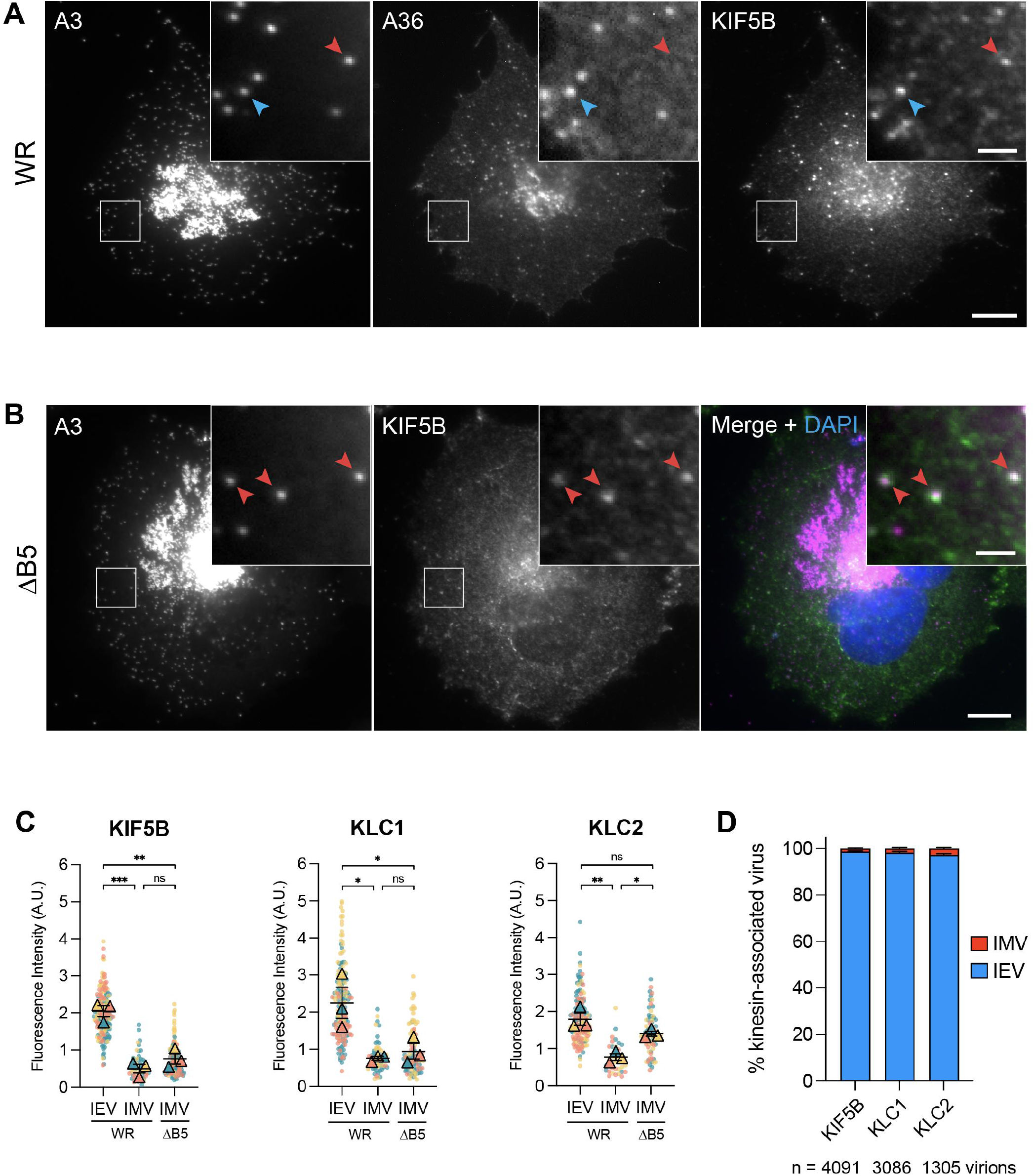
IMV and IEV associate with endogenous kinesin-1 in infected cells. **(A)** Representative immunofluorescence images of a HeLa cell labelled with A36 and KIF5B antibodies 7.5 hours post infection with WR RFP-A3. Boxed regions highlight IEV (blue arrowhead) or IMV (red arrowhead) associated with kinesin-1 (KIF5B). Scale bars, 10 μm or 2μm (inset). **(B)** Immunofluorescence images of a HeLa cell labelled with a KIF5B antibody 7.5 hours post infection with the ΔB5 RFP-A3 virus. Boxed regions highlight IMV (red arrowhead) associated with kinesin-1 (KIF5B). Scale bars, 10 μm or 2μm (inset). **(C)** SuperPlots of fluorescence intensities of KIF5B, KLC1 or KLC2 associated with IEV or IMV in WR- or ΔB5-infected HeLa cells 7.5 hours post infection. Error bars represent mean and SEM from 3 independent experiments (n = 33-174 measurements for each condition). Tukey’s multiple comparison test was used to determine statistical significance; ns, p > 0.05, * p ≤ 0.05, ** p ≤ 0.01, *** p ≤ 0.001. **(D)** Bar graph showing the percentage of KIF5B-, KLC1, or KLC2-associated virions in WR-infected cells that are either IEV or IMV. Error bars represent mean and SEM from 3 independent experiments, n = 1305-4091 kinesin-associated virions from 32-35 cells.

A36 recruits kinesin-1 to IEV by interacting with the KLC tetratricopeptide repeats (TPR) (Dodding et al., 2011; Gao et al., 2017; Rietdorf et al., 2001; Ward and Moss, 2004). We found that IMV also recruit the KLC TPR domain, suggesting they recruit kinesin-1 via a similar mechanism to IEV (Fig. S2A and B). However, examination of the sequence of IMV surface proteins fails to identify obvious W-acidic or Y-acidic motifs that mediate interactions with TPR repeats (Dodding et al., 2011; Pernigo et al., 2018; Pernigo et al., 2013; Yip et al., 2016). A lack of these motifs and a different interaction with the TPR domain such as that seen for JIP3 (Cockburn et al., 2018) may explain why during WR-infection, 97-99% of all virions with kinesin-1 were IEV (Fig. 3D). This result is consistent to our in vitro observations using extracts from A36-YdF-YFP RFP-A3 infected cells, where IEV accounted for ~98% of all virus runs (Fig. S2C). Moreover, the low incidence of kinesin-1 association with IMV likely explains why this connection was missed in previous studies.

The IMV membrane protein, A27, has been implicated in virion transport as its loss (using a virus with inducible A27 expression) resulted in an absence of IMV dispersion away from their perinuclear site of assembly (Sanderson et al., 2000). However, there is conflicting evidence, as IMV are still motile in cells infected when the A27 gene is deleted (Ward, 2005). To investigate if A27 is required for IMV transport we performed in vitro motility assays using the ΔA27 virus, which like the ΔB5 strain, only produces IMV (Ward, 2005). We found that loss of A27 has no impact on virion motility, indicating that A27 is not essential for IMV transport in vitro (Fig. S2 D and E).

### Kinesin-1 drives microtubule dependent movement of IMV and IEV

To explore the involvement of kinesin-1 in IMV motility, we infected a kinesin-1 knockout (KO) HeLa cell line generated by CRISPR/Cas9 targeting of the KIF5B gene (Jia et al., 2017) as well as a KIF5B rescued line stably expressing TagGFP2-KIF5B (Fig. S3A). To assess the role of kinesin-1 in IMV transport we quantified the total number and proportion of IMV that reached within 5 μm of the cell periphery in cells with or without KIF5B (Fig. 4A). This direct comparison is possible as there is no significant difference in cell size in the presence or absence of KIF5B (Fig. S3B). Our analysis reveals that IMV spread was impaired in the absence of KIF5B as significantly fewer IMV reached the cell periphery (Fig. 4, B and C), despite the ability of IMV to slowly disperse by random diffusion (Sodeik, 2000). Importantly, this defect was rescued by the stable expression of TagGFP2-KIF5B in the KIF5B KO cell line (Fig. 4, B and C). In parallel, we also analysed the impact of the loss of kinesin-1 on IEV transport to the cell periphery using the A36-YdF RFP-A3 virus, which is deficient in actin-based transport (Fig. 4D). We found that in the absence of kinesin-1 there was a dramatic reduction in the percentage of cells with peripheral IEV accumulations, that could be rescued by expression of TagGFP2-KIF5B (Fig. 4, E and F). Furthermore, TagGFP2-KIF5B co-localised with IEV at the cell periphery in the KIF5B rescued cell line (Fig. 4E).

**Figure 4.**
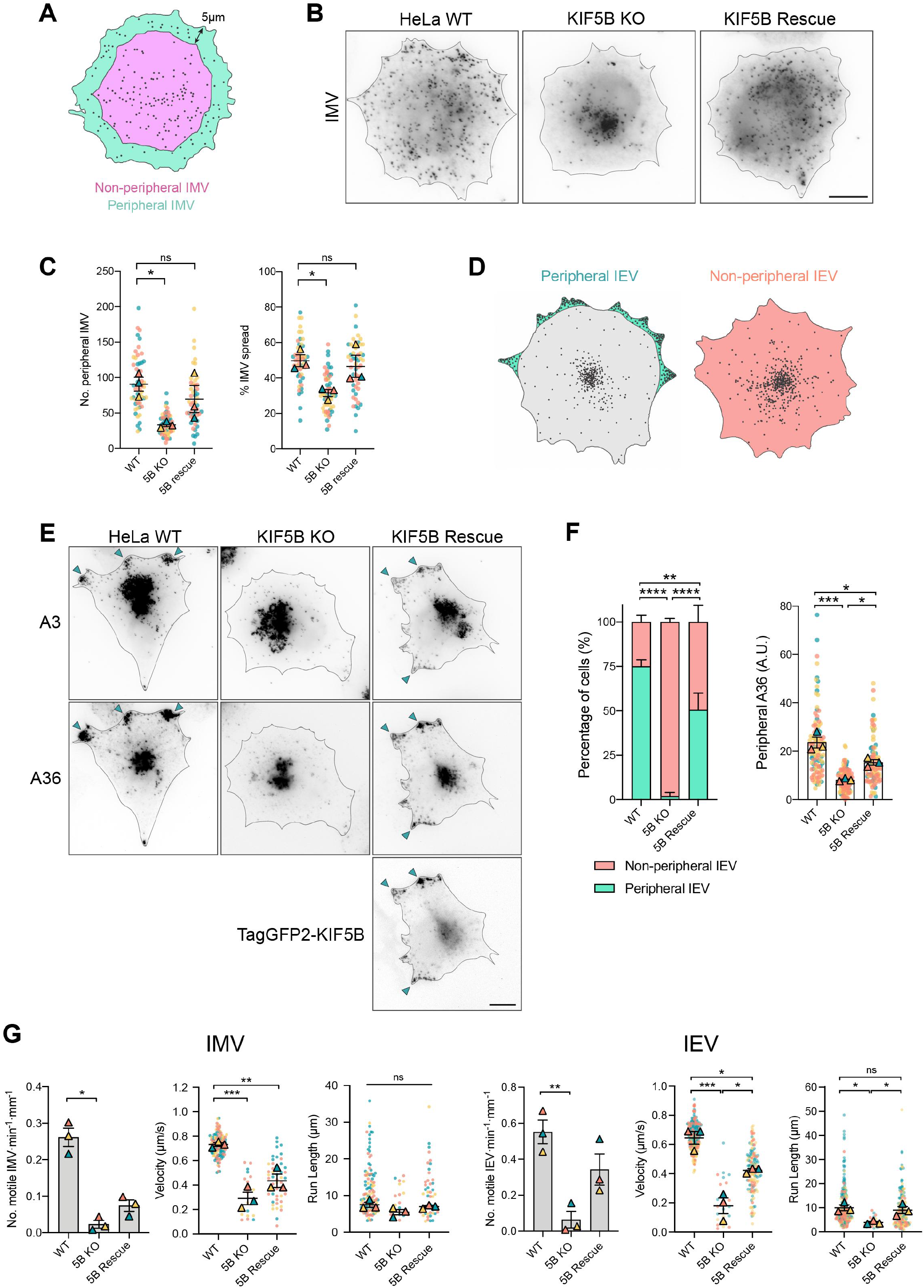
Loss of kinesin-1 impairs IMV and IEV spread and motility. **(A)** Schematic illustrating the area corresponding to the peripheral region < 5 μm from the cell edge (teal) and non-peripheral area (pink) > 5 μm from the cell edge. IMV within each region were counted to determine the total number and proportion of IMV reaching the cell periphery 7.5 hours post infection. **(B)** Representative inverted immunofluorescence images showing dispersion of IMV, labelled with an antibody detecting the IMV membrane protein A27, in indicated cell lines at 7.5 hours post infection with ΔB5 RFP-A3. Scale bar, 10 μm. **(C)** SuperPlots showing quantification of the number of peripheral IMV (left) and percentage of total IMV (right) at the cell periphery in indicated cell lines from > 50 cells in 3 independent experiments. Error bars represent the mean and SEM. Dunnet’s multiple comparison test was used to determine statistical significance; ns, not significant, p > 0.05, * p ≤ 0.05. **(D)** Illustration showing the accumulation of IEV at the perinuclear region and cell vertices (shaded green left cell) or lack of accumulation at the cell vertices (shaded pink, right cell). **(E)** Representative inverted immunofluorescence images labelled with the indicated markers showing IEV spread in indicated cell lines 7.5 hours post infection with WR A36-YdF RFP-A3 virus and labelled with A36 antibody. Scale bar, 10 μm. **(F)** Bar charts showing the percentage of cells with peripheral IEV accumulation (left) and quantification of IEV spread to the cell periphery based on fluorescence intensity of A36 antibody. Error bars represent mean and SEM from > 50 cells in 3 independent experiments. Tukey’s multiple comparison test was used to determine statistical significance; ns, p > 0.05, * p ≤ 0.05, ** p ≤ 0.01, *** p ≤ 0.001, **** p ≤ 0.0001. **(G)** SuperPlots of the in vitro motility rate, velocity and run lengths for IMV (n = 116, 18 or 48) and IEV (n = 227, 23 or 124) virions in extracts of the indicated infected cells respectively. Error bars represent mean and SEM from 3 independent experiments. Tukey’s multiple comparison test was used to determine statistical significance; ns, p > 0.05, * p ≤ 0.05, ** p ≤ 0.01, *** p ≤ 0.001.

To extend these observations, we also assessed IMV and IEV motility in vitro using extracts from infected parental, KIF5B KO or KIF5B rescue HeLa cells. Strikingly, there was a 91% reduction in microtubule-based transport of IMV in extracts lacking KIF5B compared to the parental HeLa control (Fig. 4G). In the absence of kinesin-1, the few motile IMV had a 2.6-fold reduction in velocity, 0.28 ± 0.05 μm/s compared to 0.73 ± 0.01 μm/s for the control (Fig. 4G). The partial recovery of this phenotype in the KIF5B rescue cells may be due to the reduced concentration of kinesin-1 in the extract because of the low expression of TagGFP2-KIF5B in the rescued cells (Fig. S3A). Similarly, IEV displayed negligible rates of motility and reduced velocities/run lengths in the absence of KIF5B that were also partially rescued by the presence of TagGFP2-KIF5B in extracts from A36-YdF-YFP RFP-A3 infected cells (Fig. 4G). Taken together, our observations demonstrate that kinesin-1 mediates the spread of both IMV and IEV from their perinuclear site of assembly to the cell periphery.

Kinesin-1 is clearly the major motor driving IMV and IEV motility. However, in the absence of kinesin-1, limited numbers of IMV and IEV are still weakly processive in vitro, suggesting that they may utilise additional kinesin member(s) for microtubule-based motility (Fig. 4G). The recruitment of multiple motor classes may help virions navigate the heterogenous microtubule network of the cell as different kinesins have preferences for specific microtubule subsets marked by their post-translational modifications and/or microtubule-associated proteins (MAPs). This has been well documented in neuronal cells (Hammond et al., 2010; Lipka et al., 2016) but has also been observed in non-neuronal cell types (Cai et al., 2009; Guardia et al., 2016). In line with this, kinesin-1 (KIF5B) and kinesin-3 (KIF13B) motors drive efficient transport of Rab6-positive vesicles along different microtubule populations to reach the cell periphery where they undergo exocytosis (Serra-Marques et al., 2020). In future studies, it will be interesting to resolve whether other kinesin members are also recruited by vaccinia virus and if they cooperate with kinesin-1 to promote virion transport.

### IEV and IMV recruit large but differing numbers of kinesin-1 motors

Our immunofluorescence analysis indicates that IMV and IEV recruit different numbers of kinesin-1 motors (Fig. 3C). Given that the absolute number of kinesin-1 motors on a bona fide cellular cargo remains to be established, we set out to determine the absolute number of kinesin-1 complexes recruited to IMV and IEV. To achieve this, we used a similar approach as Akamatsu et al. (2020) using a calibration curve derived from imaging nanocages with a known number of attached GFP molecules that are tethered to the plasma membrane (Fig. 5A). To extend the previous calibration curve, we generated an additional nanocage with 180 TagGFP2 molecules, then measured its background-subtracted fluorescence intensity together with the previously described 24-, 60- and 120-mer nanocages using spinning disk confocal microscopy (Fig. 5B). The average fluorescence intensity values were proportional to the predicted numbers of TagGFP2 per nanocage, including the new 180 species (Fig. 5C).

**Figure 5.**
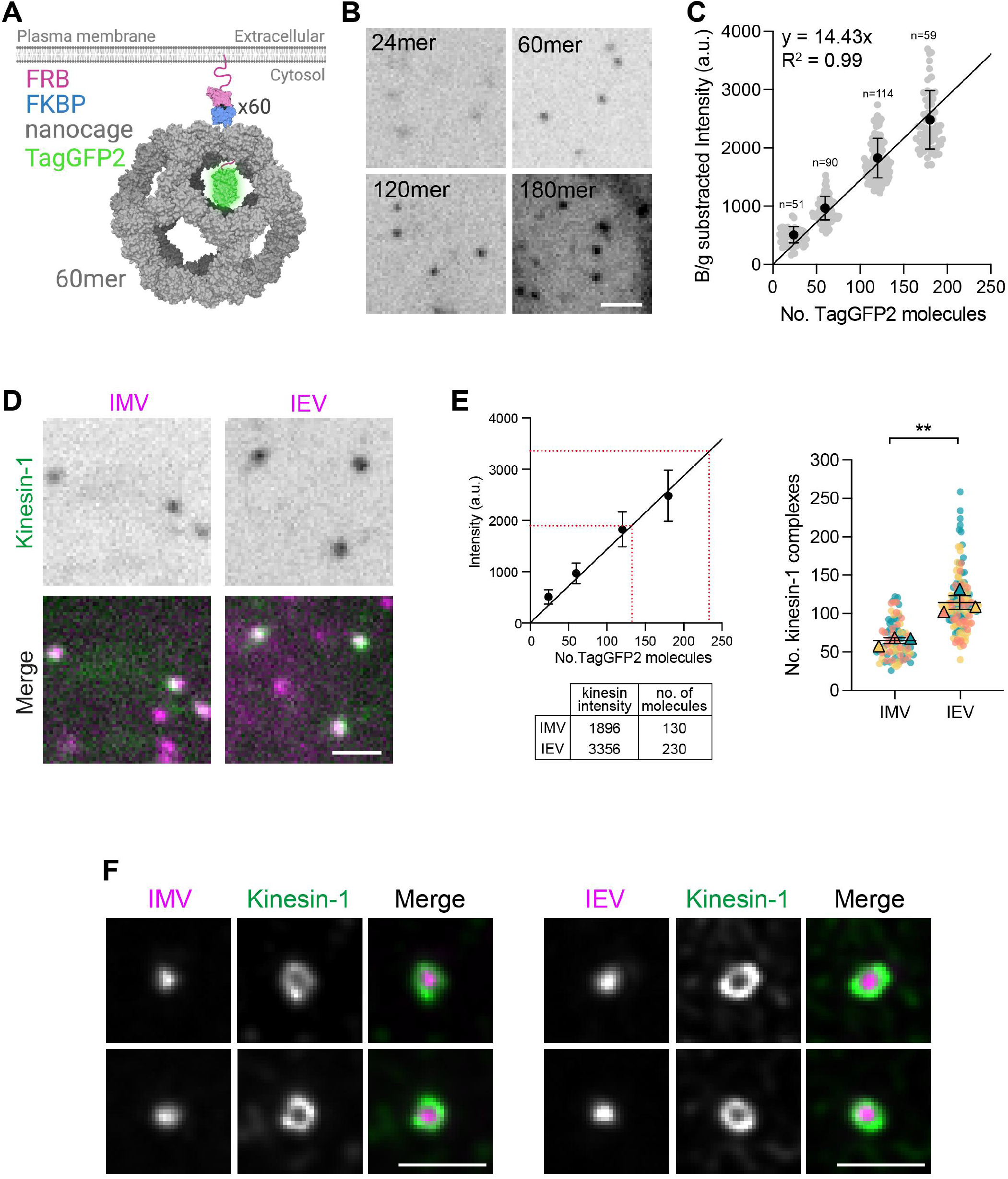
Quantifying the number of kinesin-1 complexes on IMV and IEV. **(A)** Cartoon of the intracellular TagGFP2-tagged 60mer nanocage. Each subunit of the nanocage (grey) is fused with TagGFP2 (green) and FKBP (blue), although this is shown for one subunit only for clarity. FRB (pink) is targeted to the plasma membrane by a palmitoylation and myristoylation sequence and dimerises with FKBP in the presence of rapamycin analogue AP21967. PDB structures used: 5kp9, 2y0g, 4dri. **(B)** Representative average intensity projection images of transiently expressed TagGFP2-tagged nanocages in HeLa cells treated with 500 nM AP21967. Scale bar, 2 μm. (**C**) Quantification of fluorescence intensities of TagGFP2-tagged 24-, 60-, 120- and 180-mer nanocages. Error bars represent mean and SD. Linear line of regression is fitted. N = 51-114 measurements per nanocage from 3 independent experiments. (**D**) Representative average intensity projections of endogenously expressed TagGFP2-KIF5B (green) on IEV or IMV (magenta) in HeLa TagGFP2-KIF5B knock-in cells 7.5 hours post infection with ΔB5 RFP-A3 (left) or WR B5-mRFP (right). Scale bar, 2 μm. **(E)** (Left) The average fluorescent intensity of TagGFP2-KIF5B on IMV and IEV are superimposed on the TagGFP2-tagged nanocage calibration plot from C together with the calculated number of molecules. (Right) A SuperPlot of the number of kinesin-1 complexes associated with IMV or IEV from 3 independent experiments in which 84 and 121 virions were analysed for IMV and IEV respectively. Bars represent mean and SEM. Student’s T-test was used to determine statistical significance; ** p ≤ 0.01. **(F)** Representative maximum intensity projections of deconvolved structured illumination images showing the localisation of kinesin-1 (green), detected with the KLC1 antibody on IMV and IEV (both magenta) at 7.5 hours post infection with ΔB5 RFP-A3 (left) or WR B5-mRFP (right). Scale bars, 1 μm.

To compare the fluorescence intensities of TagGFP2-tagged nanocages with kinesin-1 associated with IMV or IEV, we generated an endogenously expressed TagGFP2-KIF5B HeLa knock-in cell line by CRISPR/Cas9 genome editing and single cell cloning (Fig. S3C). As with our previous TagGFP2-KIF5B rescue cell line, immunoblot analysis showed that TagGFP2-KIF5B expression is reduced in the knock-in cell line compared to the untagged motor in the parental cells (Fig. S3C). Nevertheless, all kinesin-1 motors in the knock-in cell line are fluorescently tagged and amenable for analysis. As the molecule counting method requires imaging z-stacks, the fast microtubule-based movements of IMV and IEV presents a challenge for capturing the intensity of TagGFP2-KIF5B on moving virions due to the temporal constraints. Live cell imaging, however, reveals that the ratiometric intensity of viral RFP-A3 and TagGFP2-KIF5B signals in a single z-plane does not significantly change between phases of IMV motility and pausing (Fig. S3D and Video 8). We therefore quantified the number of kinesin-1 molecules on stationary IMV and IEV particles in cells infected with ΔB5 RFP-A3 or WR B5-RFP respectively. In agreement with our previous immunofluorescence analysis, we found IEV recruit more kinesin-1 than IMV (Fig. 5, D and E). Comparison of the fluorescence intensity of TagGFP2-KIF5B on each virion with our nanocage calibration curve reveals that IMV and IEV recruit an average of 130 ± 8 and 230 ± 18 (mean ± SEM) KIF5B molecules, which is equivalent to 65 ± 4 and 115 ± 9 kinesin-1 motor complexes respectively (Fig. 5E). This significant difference in motor number may explain why IEV are more efficient (longer run lengths) than IMV in their transport to the plasma membrane. Indeed, in previous experimental and theoretical studies, increasing the number of kinesin motors attached to a cargo leads to further travel distances along the microtubule (Beeg et al., 2008; Derr et al., 2012; Furuta et al., 2013; Korn et al., 2009; Muller et al., 2010; Vershinin et al., 2007). We have also observed that impairing the ability of IEV to recruit kinesin-1 by mutating one of the A36 tryptophan acidic motifs also leads to a significant reduction in run length without affecting viral speed (Dodding et al., 2011). Our motor number values are significantly larger than previous studies that typically observe 1-11 kinesin motors on a cargo using immunogold labelling in EM sections or are based on inferences from cargo velocities or force measurements (Ashkin et al., 1990; Gross et al., 2007; Hill et al., 2004; Kural et al., 2005). Due to the relatively large ≈360 × 270 × 250 nm size of IMV (Cyrklaff et al., 2005), it is perhaps not surprising that so many kinesin-1 motors are associated with each virion. Indeed, our values are more in line with the predicted number of kinesin-1 motor complexes required to transport a vesicle of similar size to a virion over long distances (>10μm) (Jiang et al., 2019).

It has been suggested that kinesin motors are likely to be clustered on cellular cargoes to ensure more efficient processive transport (Erickson et al., 2011). Given this, we wondered how the motors were spatially organised on the virion surface given the relatively large numbers of kinesin-1 on both IMV and IEV. Super-resolution imaging of fixed ΔB5- and WR-infected cells using structured illumination microscopy reveals that kinesin-1 is distributed over the whole IMV or IEV surface (Fig. 5F, S3E). Such an organisation may help the virions navigate the dense cellular microtubule network by allowing them to quickly switch microtubule tracks and/or bypass roadblocks for efficient transport in the crowded cytosol (Lakadamyali, 2014; Tjioe et al., 2019). Furthermore, given this organisation, it is likely that only a subset of bound motors is active at any given time due to the geometric constraints of motor positioning on the virion relative to a microtubule. Indeed, it is predicted that for a 100 nm vesicle with 35 bound kinesin-1 motors, three motors are sufficient to engage the microtubule and drive vesicle transport over long 10 μm distances (Jiang et al., 2019). However, determining the number of active motors on a cargo in live-cells still remains a considerable challenge (Cai et al., 2007), as while cargo binding relieves auto-inhibition it may not always result in full motor activation (Blasius et al., 2007; Fu and Holzbaur, 2013; Kawano et al., 2012; Twelvetrees et al., 2019) without additional regulation through post-translational modifications and/or microtubule-associated proteins (Chiba et al., 2021; Hooikaas et al., 2019; Manser et al., 2012). Given our observations, we suggest that vaccinia infected cells offer a powerful model system with which to develop and test sensors for the activation state of kinesin-1 on moving cellular cargoes.

In conclusion, our study shows that kinesin-1 drives the transport and spread of both intracellular forms of vaccinia virus. In addition, we show for the first time that microtubule-based motility of both IMV and IEV can be reconstituted in infected cell extracts in vitro. This will no doubt provide a useful model system to obtain further insights into motor-cargo relationships and motor regulation. The task ahead is to uncover the mechanistic basis for kinesin-1 recruitment to IMV and determine how kinesin-1 recruitment and activation is regulated by IMV and IEV.

## Acknowledgements

We would like to thank Gil Henkin, Tanja Consolati, and Jonathon Hannabuss from the Thomas Surrey lab (Francis Crick Institute, UK) for their help and expertise in establishing the in vitro microtubule motility assays as well as Todd Fallesen (Crick CALM STP) for providing the protocol for polarity marked MTs. We also acknowledge Drs Juan Bonifacino (NIH, USA), Geoffrey Smith (University of Cambridge, UK), Brian Ward, University of Rochester Medical Center, New York, USA), Marvin Bentley (Rensselaer Polytechnic Institute, New York, USA) and David Drubin (UC Berkeley, USA) for providing published reagents. We thank Matthew Akamatsu (Drubin lab) for his input during optimisation of nanocage imaging, Snezhka Oliferenko and Jeremy Carlton (the Francis Crick Institute) for helpful comments on the manuscript and the Way lab for scientific discussions during the project. For the purpose of Open Access, the authors have applied a CC BY public copyright license to any Author Accepted Manuscript version arising from this submission.

## Author contributions

The generation of all stable cell lines, experiments and data analysis was performed by A. Xu. The A36-YdF and A36-YdF-YFP viruses containing RFP-A3 as well as WR B5-RFP were generated by T.P. Newsome and the ΔB5 RFP-A3 virus was constructed by S. Schleich. A. Basant generated the 180mer TagGFP2 nano cage and optimised conditions for nanocage imaging and analysis. M. Way conceived and supervised the work with the help of A. Basant. A. Xu and M. Way wrote the manuscript with input from the other authors.

## Materials and Methods

### Cells and generation of stable cell lines

HeLa cells lines were maintained in minimal essential medium (MEM) or Dulbecco’s modified eagle medium (DMEM) supplemented with 10% FBS, 100 U/ml penicillin, and 100 μg/ml streptomycin at 37°C and 5% CO_2_. Stable HeLa cell lines expressing GFP-KLC1 and GFP-KLC2 were generated using the lentivirus infection (Trono group second generation packaging system, Addgene) and selected using puromycin resistance (1 μg/ml) as previously described (Abella et al., 2016). The HeLa KIF5B KO cell line was kindly provided by Juan Bonifacino (NIH, USA) (Jia et al., 2017). Lentiviral expression vectors were used to stably express tagGFP2-KIF5B in HeLa KIF5B KO cells to generate the HeLa KIF5B rescue cell line.

The vectors pLVX GFP-KLC1 and pLVX GFP-KLC2 were generated by sub-cloning the murine KLC1A and KLC2 coding sequences (Dodding et al., 2011) into the EcoRI/BamHI and NotI/EcoRI sites respectively of a pLVX N-term-GFP parent vector (Abella et al., 2016). To generate the pLVX tagGFP2-KIF5B vector, the murine KIF5B coding sequence was amplified from a plasmid provided by Marvin Bentley (Rensselaer Polytechnic Institute, New York, USA) (Yang et al., 2019), tagGFP2 was amplified from a plasmid provided by David Drubin (UC Berkeley, USA) (Akamatsu et al., 2020) and both inserted between the XhoI and EcoRI sites of the parental pLVX N-term GFP vector using Gibson Assembly (New England Biolabs), following the manufacturer’s instructions. These lentivirus vectors were used to establish stable HeLa cell lines as previously described (Weisswange et al., 2009). SnapGene software (Insightful Science; available at snapgene.com) was used to plan and visualise cloning strategies, and to analyse sequencing results.

### Expression constructs

The expression vector pEL KLC2-TPR has been described previously (Rietdorf et al., 2001), KLC sequences comprising residues 1-155 of murine KLC2 and residues 1-162 or 163-538 of murine KLC1A were amplified by PCR and cloned into the NotI/EcoRI site of the pEL N-term GFP parental vector using Gibson Assembly (New England Biolabs) following the manufacturer’s instructions (Rietdorf et al., 2001). The KLC1A and KLC2 coding sequences used for PCR amplification have been previously described (Dodding et al., 2011). The fidelity of all expression constructs was confirmed by sequencing.

### CRISPR/Cas9 mediated gene editing

The HeLa CRISPR-Cas9 knock-in cell line expressing TagGFP2-KIF5B at the endogenous KIF5B locus was generated using the pORANGE vector containing SpCas9, purchased from Addgene (Plasmid #131471) (Willems et al., 2020). The guide RNA (gRNA) for KIF5B was designed using a CRISPR design webpage tool (https://www.benchling.com/). The targeting sequence used was as follows (coding strand sequence indicated): 5′-CCCGGCTGCGAGAAAGATGG-3’. CRISPR/Cas9 mediated knock-in of TagGFP2 into the endogenous KIF5B locus was performed according to the protocol described in Willems et al. (2020). In brief, HeLa cells were transfected using JetPrime (Polyplus) with the pORANGE vector bearing the appropriate gRNA targeting sequence and TagGFP2 insert. The gRNA targets the ATG start codon of KIF5B exon 1 where Cas9 induces a double strand break. The TagGFP2 coding sequence is integrated into the incision site through repair by non-homologous end joining (NHEJ). After initial transfection, cells were allowed to recover for ~3 weeks before single-cell colonies were isolated by FACS. Individual clones were screened for biallelic integration of TagGFP2 into the KIF5B loci by junction PCR and immunoblot analyses. Sequencing confirmed successful in-frame integration of the TagGFP2 sequence.

### Recombinant viruses and infection

All recombinant viruses are generated in the Western Reserve (WR) strain of Vaccinia virus. The recombinant vaccinia virus strains RFP-A3 and ΔA27 YFP-A4 (kindly provided by Brian Ward, University of Rochester Medical Center, New York, USA) have previously been described (Ward, 2005; Weisswange et al., 2009). The LA-RFP-A3-RA targeting vector was used to insert RFP-A3 as previously described (Weisswange et al., 2009) into the genome of the existing viral strains ΔB5 (Engelstad and Smith, 1993), A36-YdF (Rietdorf et al., 2001), and A36-YdF-YFP (Arakawa et al., 2007). The recombinant strain expressing B5-RFP was generated by rescuing the ΔB5 virus with a B5-RFP targeting vector and selecting for an increase in plaque size to that of WR and expression of RFP.

For live and fixed cell imaging, HeLa cells on fibronectin-coated coverslips or glass bottomed dishes were infected with the relevant recombinant vaccinia virus in serum-free DMEM at a multiplicity of infection (MOI) of 1. After one hour at 37°C, the serum-free DMEM was removed and replaced with complete DMEM. Cells were incubated at 37°C and at 7.5 hours post-infection cells were imaged live or were fixed and processed for immunofluorescence analysis. For nocodazole experiments, DMSO control or 33 μM nocodazole was added to culture medium for 1 h prior to fixation or live-cell imaging.

### Immunofluorescence and immunoblot analysis

HeLa cells were fixed with 4% paraformaldehyde in PBS for 10 min, blocked in cytoskeletal buffer (10 mM MES, 150 mM NaCl, 5 mM EGTA, 5 mM MgCl_2_, and 5 mM glucose pH 6.1) containing 2% (v/v) fetal calf serum and 1% (w/v) BSA for 30 min, and then permeabilised with 0.1% Triton X-100 in PBS for 5 min. To differentiate between IMV and IEV, cells were stained with a monoclonal antibody against A36 (1:50) kindly provided by Geoffrey Smith (University of Cambridge, UK) (van Eijl et al., 2000) followed by a Cy5 goat anti-mouse secondary antibody (115-175-146; Jackson ImmunoResearch, 1:1,000). To visualise IMV in ΔB5-infected cells, a monoclonal antibody against A27 (1:1,000) was used followed by a Cy5 goat antimouse secondary antibody (Rodriguez et al., 1985). To visualise kinesin-1, the following primary antibodies were used at the specified dilutions: KIF5B (ab167429; Abcam, 1:400), KLC1 (sc-25735; Santa Cruz, 1:400), KLC2 (HPA040416; Atlas Antibodies, 1:400), followed by an Alexa Fluor 488 goat anti-rabbit secondary antibody (A11034; Invitrogen, 1:1,000). Coverslips were mounted on glass slides using Mowiol (Sigma) and image acquired on a Zeiss Axioplan2 microscope equipped with a 63x/1.4 NA Plan-Achromat objective and a Photometrics Cool Snap HQ cooled charge-coupled device camera. The microscope was controlled with MetaMorph 7.8.13.0 software. Images were analysed using Fiji and processed with Adobe software package.

For structured illumination microscopy, samples were prepared as above and imaged on an Olympus iX83 Microscope with Olympus 150x/1.45 NA X-Line Apochromatic Objective Lens, dual Photometrics BSI-Express sCMOS cameras and CoolLED pE-300 Light Source (Visitech) and was controlled using Micro-Manager 2.0.0. Image stacks of 10-15 z-slices with 0.1 μm steps were acquired and deconvolved using the express deconvolution setting on Huygens Software (Scientific Volume Imaging).

For immunoblot analyses, the following antibodies were used at indicated dilutions: β-tubulin (T7816; Sigma, 1:10,000), GFP clone 3E1 (Francis Crick Institute, Cell Services STP, 1:5,000), KIF5B (ab167429; Abcam; 1:1,000), KLC1 (sc-25735; Santa Cruz; 1:1,000), KLC2 (HPA040416; Atlas Antibodies; 1:1,000), and A27 (C3 monoclonal, 1:1,000) (Rodriguez et al., 1985).

### Live-cell imaging and Automated Particle Tracking in cells

Live-cell imaging experiments were performed at 7.5 hours post-infection in complete DMEM (10% FBS, 1 % P/S) in a temperature-controlled chamber at 37°C. Cells were imaged on a Zeiss Axio Observer microscope equipped with a Plan Achromat 63x/1.40 NA Ph3 M27 oil lens or a Plan Achromat 100x/ 1.46 NA oil lens, an Evolve 512 camera, and a Yokagawa CSUX spinning disk. The microscope was controlled by the SlideBook software (3i Intelligent Imaging Innovations). Time lapse images used for automated particle tracking were acquired at a sampling rate of 10 Hz using an exposure of 33 ms for the RFP (virus) channel.

To quantify the number of kinesin-1 associated with virions, image stacks of 10 z-slices, 0.1 μm apart were acquired at 0.2 Hz using an exposure of 100 ms or 30 ms for the respective GFP (kinesin) and RFP (virus) channels. All other movies were typically imaged at 1 Hz using an exposure of 100 ms for each channel. To visualise microtubules in live infected cells, 125 nM SiR-Tubulin (Cytoskeleton #CY-SC002) was added to the culture medium 2 h prior to imaging and were visualised using the FarRed channel.

To track viruses in infected cells, we used the Fiji plugin, TrackMate (Tinevez et al., 2017). The Laplacian of Gaussian (LoG) detector identified virion spots with an estimated diameter of 1 μm and threshold of 30 using a median filter and sub-pixel localisation. We generated whole virus trajectories using the simple linear assignment problem (LAP) tracker with a linking distance of 0.8 μm and gap closing distance of five frames. These were filtered using a track displacement threshold > 1 μm. The track data was exported and processed using TraJ (Wagner et al., 2017) to analyse the trajectories, categorising virus tracks into segments (sub-trajectories) representing either active transport, normal diffusion, sub-diffusion, or confined motion. The data for active transport was exported to Excel to derive the virion velocities and run lengths.

### In vitro virus motility assays

HeLa cells were grown in 10 cm culture plates until ~80% confluent then infected with relevant virus for 18 h at a multiplicity of infection (MOI) of 0.1. Infected cells were detached by versene treatment and centrifuged (1700 rpm, 5 min, 4°C). The cell pellet was resuspended in 1-pellet volume of assay buffer (40 mM HEPES, 1 mM EGTA, 1 mM MgCl_2_, 100 mM KCl, 1% Glucose, 1 mM GTP, 10 mM BME) supplemented with protease inhibitors (cOmplete™ Mini EDTA-free, Sigma) and 1 mM phenyl-methyl-sulfonyl fluoride (PMSF). Cells were lysed through two iterative freeze/thaw cycles and the cell lysate was clarified by centrifugation (1700 rpm, 5 min, 4°C). The cell extract was kept on ice for not more than 2 h before the start of imaging. An ATP regeneration system (2 mM ATP, 25 mM phosphocreatine and 0.013 mg/ml creatine phosphokinase at >150 Units/ml) and an oxygen scavenging system (12.5 mg/ml glucose oxidase and 3 mg/ml catalase) was added to the extract prior to adding into the flow chamber.

Tubulin was purified from pig brains as previously described (Castoldi and Popov, 2003). GMPCPP-stabilised microtubules were polymerised from unlabelled tubulin (1.42 μM), biotin-labelled tubulin (0.5 μM) and AlexaFluor 647-labelled tubulin (0.27 μM) in 1X BRB80 (80 mM K-PIPES pH 6.8, 1 mM MgCl2, 1 mM EGTA pH 6.8) containing GMPCPP (0.5 μM) for 3 h at 37°C. Polymerised microtubules were pelleted by centrifugation at 11,500 rpm (12,420 rcf) for 5 min, gently resuspended in BRB80 and left in dark at room temperature overnight for use the following day. Polarity-marked microtubules stabilised with taxol were generated as previously described (Fallesen et al., 2017). Glass coverslips were functionalized with a layer of biotin and biotin-PEG (Rapp Polymere), while glass slides were passivated with PLL-PEG (SuSoS) as previously described (Bieling et al., 2010). Flow chambers forming a ~10 μL volume chamber (chamber size ~0.5 × 18 × 0.1 mm), consisted of a biotin-PEG functionalised coverslip attached to a PLL-PEG passivated glass slide via double-sided tape (Tesa, Hamburg). The glass surfaces were passivated with 5% Pluronic F-127 (Sigma) followed by κ-casein (0.05 mg/ml) for 10 min each, then incubated with NeutrAvidin (Invitrogen) for 3 min. Polymerised microtubule mix was then added for 10 min before unattached microtubules were removed through several washes using assay buffer. Finally, infected cell extract in assay buffer was added and chambers were sealed with Vaseline (Unilever) prior to imaging on a spinning-disc microscope at 37°C. Images were acquired at 1 frame per second, and each sample was imaged for no longer than 30 min.

### Image analysis and quantitation

Fluorescence intensity of kinesin-1 antibody labelling was analysed in fixed-cell images using Fiji (Schindelin et al., 2012). Raw integrated kinesin-1 antibody signal was measured following the approach of (Verdaasdonk et al., 2014) by drawing an eight-pixel circle over kinesin-1 spots colocalising with a virion. The background signal was obtained by drawing a larger ten-pixel concentric circle and measuring the raw integrated density. The area-corrected background intensity was subtracted from the initial eight-pixel region of interest to acquire the fluorescence intensity per kinesin-1 spot.

The constructs that assembled into 24-, 60- and 120-mer nanocages used for the fluorescence calibration standard curve were generated as described previously (Akamatsu et al., 2020). To generate a construct that would self-assemble into 180mers, plasmids obtained from the Drubin lab were modified (Akamatsu et al., 2020). The NheI/XbaI fragment was replaced by a synthetic construct (Invitrogen; Geneart) where the KPDG aldolase was tagged at the N terminus with two TagGFP2. GS repeat linkers were included between TagGFP2 sequences. The nanocage constructs were transiently expressed for ~26 h in HeLa cells after transfecting cells with Lipofectamine 2000 (Invitrogen) prior to adding 500 nM AP21967 (Takara) to the medium 30 min before imaging to induce self-assembly of nanocages. Average intensity projections of image z-stacks were used to measure the fluorescence intensity per nanocage spot. The background-subtracted intensity of each nanocage was measured as described above and plotted as a function of predicted TagGFP2 copy number per nanocage to obtain the calibration standard curve. A line of linear fit through the origin was applied by linear least-squares fitting. Identical analysis was performed on kinesin-1 spots that colocalised with virions in infected cells to calculate the number of kinesin-1 associated with IMV or IEV. Quantification was only performed on single stationary spots.

To quantify the number of motile IMV in nocodazole experiments, TrackMate was used to identify virus trajectories. A displacement threshold was applied to all tracks to quantify the number of viruses that moved > 3 μm within the 1 min imaging window.

In vitro virus motility was analysed by kymograph analysis using the ImageJ Kymograph plugin made by Jens Rietdorf and Arne Seitz (EPFL Lausanne). The Fiji line tool was used to measure constant velocity segments within kymographs and the data was exported to Excel to derive the virus velocities and run lengths. Virus motility rates were calculated as the total number of motile virions detected, normalised to the imaging duration (in minutes) and microtubule length (in mm) within each field of view. The overall virus motility rate per independent experiment is reported.

## Supplementary figure legends

**Supplementary Figure 1.**
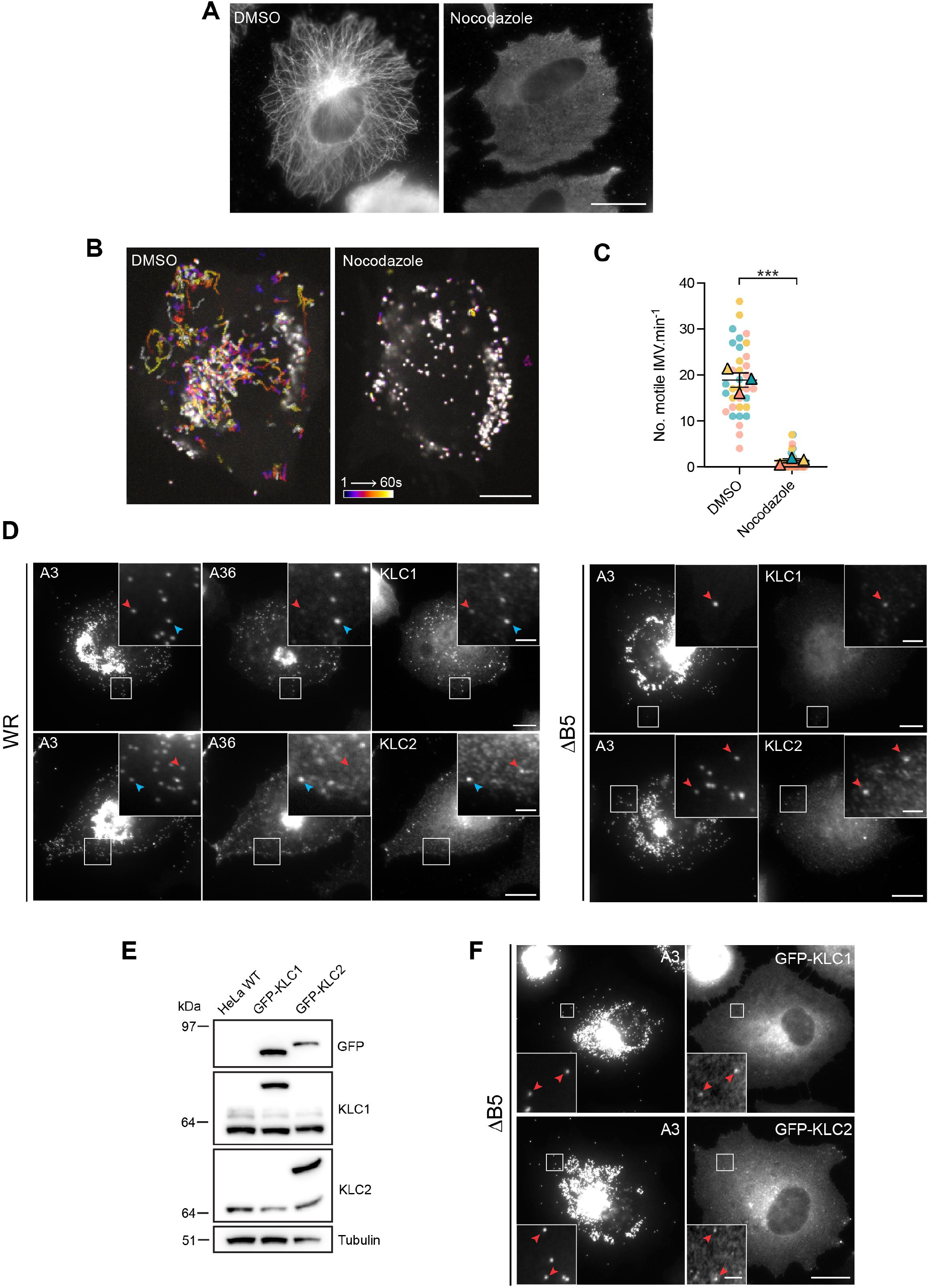
IMV recruit kinesin-1 to undergo microtubule-based motility. **(A)** Representative immunofluorescence images showing the organization of microtubules using a tubulin antibody in HeLa cells infected with the ΔB5 virus for 7.5 h and treated with DMSO or 33 μM nocodazole for 1 h. Scale bar, 10 μm. **(B)** Representative maximum intensity projection images showing the movement of IMV in HeLa cells infected with ΔB5 RFP-A3 for 7 h and treated with DMSO or 33 μM nocodazole for 1 h prior to imaging for 60 s as indicated by the timestamp bar (see Video 3). Scale bar, 10 μm. **(C)** SuperPlot quantifying the number of motile IMV (defined as IMV travelling > 3 μm) during the 60 s imaging window in infected cell treated with DMSO or 33 μM nocodazole for 1 hour. n = 34 cells per condition from 3 independent experiments. Student’s T-test was used to determine statistical significance; *** p ≤ 0.001. **(D)** Representative immunofluorescence images of HeLa cells infected with WR RFP-A3 (left panel) or ΔB5 RFP-A3 (right panel) labelled with either KLC1 or KLC2 and A36 antibodies. Boxed regions highlight IMV (red arrowheads) and IEV (blue arrowheads) associated with endogenous KLC1 or KLC2. Scale bars, 10 μm and 2 μm (inset). **(E)** Immunoblot analyses with the indicated antibodies of total cell lysates from parental HeLa wild-type (WT) or HeLa cells stably expressing of GFP-KLC1 or GFP-KLC2. **(F)** Representative immunofluorescence images showing the recruitment of GFP-KLC1 (top) or GFP-KLC2 (bottom) to IMV in HeLa cells stably expressing the indicated GFP-tagged protein and infected with the ΔB5 RFP-A3 virus. Red arrowheads highlight IMV colocalisation with KLC. Scale bars, 10 μm and 2 μm (inset).

**Supplementary Figure 2.**
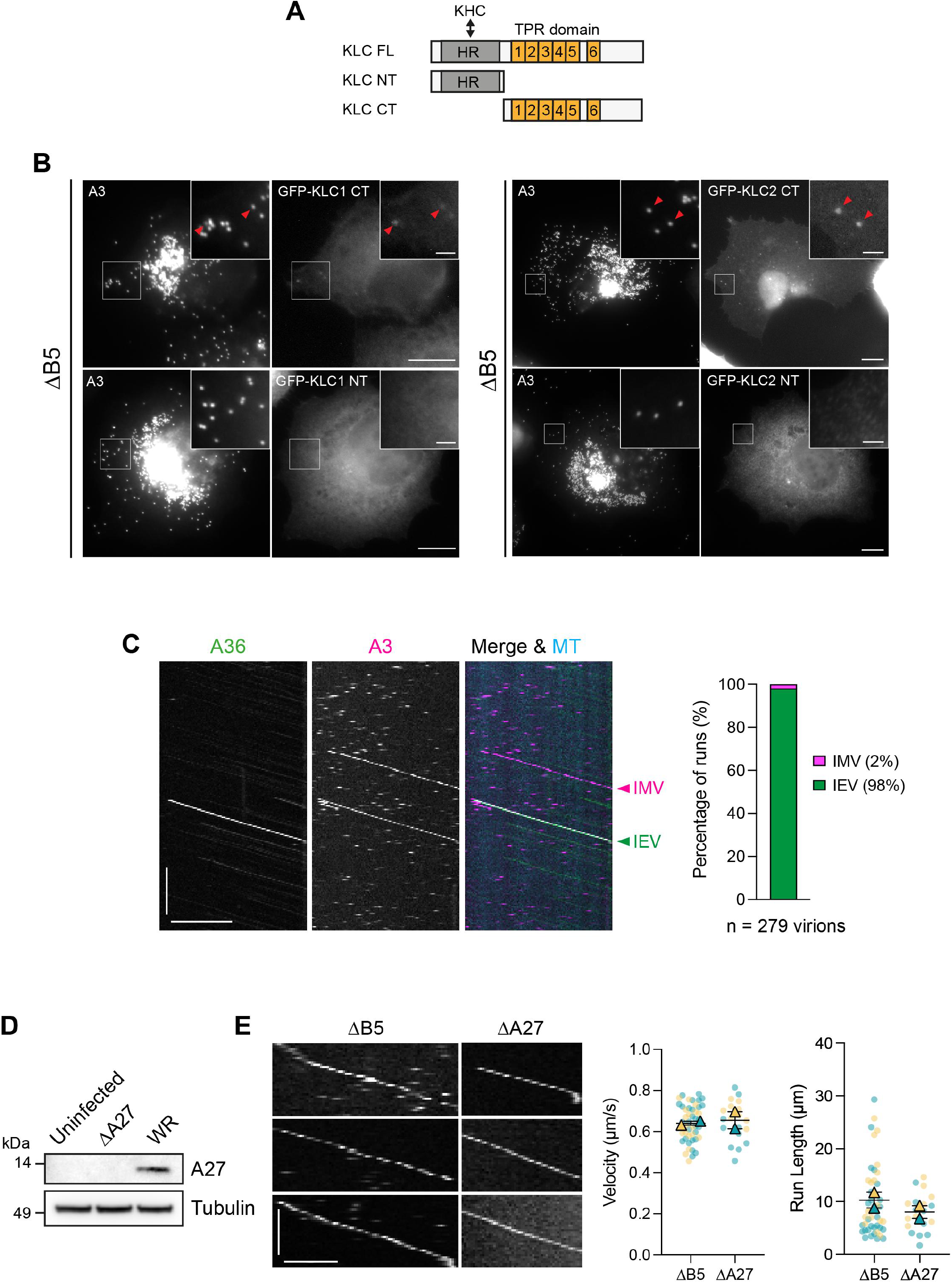
KLC C-terminal domains are sufficient for recruitment to IMV. **(A)** Schematic of KLC full length (FL), N-terminal (NT) and C-terminal (CT) constructs. The NT contains the heptad repeat (HR) domain which binds kinesin heavy chain (KHC) while the CT contains the tetratricopeptide repeat (TPR) domain involved in cargo binding. **(B)** Representative immunofluorescence images showing IMV recruits the C-terminus (CT) of GFP-tagged KLC1 and KLC2 (indicated by red arrowheads) but not their N-terminal (NT) domain in HeLa cells transiently expressing the indicated GFP-tagged protein and infected with ΔB5 RFP-A3 for 7.5 hours. Scale bars, 10 μm and 2 μm (inset). **(C)** Kymograph showing movement of IMV and IEV along the same microtubule. Scale bars, 30 s (vertical) and 10 μm (horizontal). Bar graph (right) shows the percentage of motile IMV and IEV in vitro assays using extracts from cells infected with WR A36-YdF-YFP RFP-A3. N = 279 virions from 3 independent experiments. **(D)** Immunoblot of whole cell lysates from uninfected, ΔA27- or WR-infected HeLa cells using indicated antibodies. **(E)** Example kymographs showing in vitro IMV motility in extracts derived from cells infected with ΔB5 RFP-A3 (left) or ΔA27 YFP-A4 (right) viruses. The velocities and run lengths of IMV produced by these two viruses are shown in the corresponding SuperPlots. n = 46 (ΔB5) or 19 (ΔA27) virions from 2 independent experiments.

**Supplementary Figure 3.**
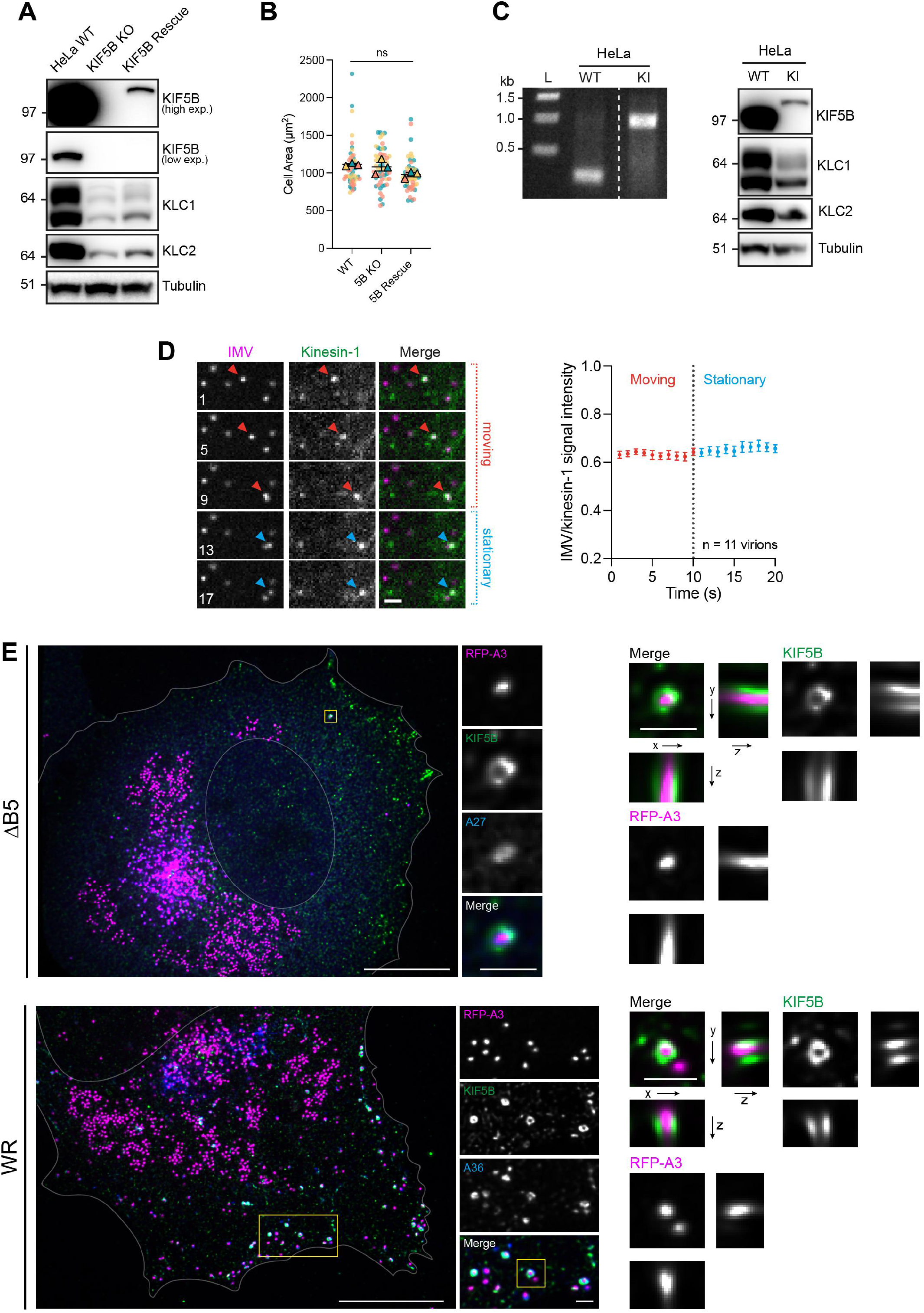
Validation of TagGFP2-KIF5B HeLa CRISPR knock-in cell line. **(A)** Immunoblot analyses with the indicated antibodies of total cell lysates from parental HeLa wild-type (WT), KIF5B knockout (KO) and KIF5B KO stably expressing TagGFP2-KIF5B (‘KIF5B Rescue ‘). **(B)** Quantification of cell area of indicated cell lines. At least 50 cells were measured from 3 independent experiments. Error bars represent mean and SEM. Ordinary one-way ANOVA test was used to determine statistical significance; ns, p > 0.05. **(C)** Agarose gel electrophoresis (left) showing PCR bands of KIF5B locus in HeLa wild-type (WT) or TagGFP2-KIF5B CRISPR knock-in (KI) cells. Immunoblot analysis (right) of total cell lysates from HeLa wild-type (WT) or TagGFP2-KIF5B CRISPR knockin (KI) cells using the indicated antibodies. **(D)** Movie stills showing the association of TagGFP2-KIF5B (green) with IMV (magenta) while moving or stationary as indicated by the red and blue arrowheads respectively in the HeLa TagGFP2-KIF5B knock-in cell line at 7.5 hours post infection with the ΔB5 RFP-A3 virus (see Video 8). Time in seconds is indicated in each image. Scale bar, 2 μm. Graph on right shows quantification of ratio metric fluorescence intensity of TagGFP2-KIF5B and RFP-A3 signals during the two phases. n = 11 virions from 2 independent experiments. **(E)** Maximum intensity projections of deconvolved super-resolution images of a HeLa cell infected with ΔB5 RFP-A3 (top) or WR RFP-A3 (bottom) and immunolabelled with KIF5B (green) and A27 (top panel, blue) or A36 (bottom panel, blue). Scale bars, 10 μm, 1 μm (insets).

## Video Legends

**Video 1. Representative example of IMV movement in an ΔB5 infected HeLa cell (also see Fig. 1A).** HeLa cell infected with ΔB5 RFP-A3 for 7.5 h before imaging. The time in minutes:seconds is indicated, and the scale bar = 10 μm. Images were taken every second. Video plays at 10 frames per second.

**Video 2. Representative examples of microtubule-based IMV movements (also see Fig. 1A).** Insets from a HeLa cell infected with ΔB5 RFP-A3 for 7.5 h before imaging. Coloured trajectories show active (left), diffusive (middle) and stationary (right) virus movements. The time in minutes:seconds is indicated, and the scale bar = 2 μm. Images were taken every second. Video plays at 10 frames per second.

**Video 3. IMV movements in the presence or absence of microtubules (also see Fig. S1B).** Example of a HeLa cell infected with ΔB5 RFP-A3 for 7.5 h and treated with DMSO (left) or 33 μM nocodazole (right) for 1 h before imaging. The time in seconds is indicated, and the scale bar = 10 μm. Images were taken every 0.1 seconds. Video plays at 60 frames per second.

**Video 4. IMV motility on GMPCPP microtubules (also see Fig. 2C)**

Example of IMV labelled with RFP-A3 (indicated by the white arrowheads) moving on GMPCPP-stabilised microtubules (blue) in vitro in the presence of 2 mM ATP (left panel) but not AMPPNP (right panel). The time in minutes:seconds is indicated, and the scale bar = 5 μm. Images were taken every second. Video plays at 10 frames per second.

**Video 5. IEV motility on GMPCPP microtubules (also see Fig. 2C)**

Example of IEV labelled with RFP-A3 and A36-YdF-YFP (indicated by the white arrowheads) moving on GMPCPP-stabilised microtubules (blue) in vitro in the presence of 2 mM ATP (left panel) but not AMPPNP (right panel). The time in minutes:seconds is indicated, and the scale bar = 5 μm. Images were taken every second. Video plays at 10 frames per second.

**Video 6. IEV towards the plus end of microtubules (also see Fig. 2D)**

Example of IEV labelled with RFP-A3 and A36-YdF-YFP (indicated by the white arrowheads) translocating towards the bright microtubule plus-end in vitro in the presence of 2 mM ATP. The time in minutes:seconds is indicated, and the scale bar = 10 μm. Images were taken every second. Video plays at 20 frames per second.

**Video 7. IMV move towards the plus end of microtubules (also see Fig. 2D)**

Example of IMV labelled with RFP-A3 (indicated by the white arrowheads) translocating towards the bright microtubule plus-end in vitro in the presence of 2 mM ATP. The time in minutes:seconds is indicated, and the scale bar = 5 μm. Images were taken every second. Video plays at 20 frames per second.

**Video 8. KIF5B associates with moving and stationary IMV (also see Fig. 3D)**

Example movie of an IMV labelled with RFP-A3 (indicated by the white arrowhead) recruiting endogenously expressed tgGFP2-KIF5B in HeLa knock-in cells. The time in minutes:seconds is indicated, and the scale bar = 2 μm. Images were taken every second. Video plays at 7 frames per second.

